# Inbreeding depression by polygenic load following a severe population bottleneck

**DOI:** 10.64898/2026.06.03.729877

**Authors:** Kiran G. L. Lee, Alessandro Pinto, Sen Dong, Sergio González-Mollinedo, Chuen Z. Lee, Jon Slate, Jan Komdeur, David S. Richardson, Hannah L. Dugdale, Terry Burke

## Abstract

It is poorly understood how populations survive extreme bottlenecks despite severe inbreeding. We investigate the genomic architecture of inbreeding in the once critically endangered Seychelles warbler (*Acrocephalus sechellensis*) using 37 years of individual-based monitoring and 1.8 million SNPs. Linkage disequilibrium patterns reveal a historical effective population size of ∼270 that plummeted to ∼13 coinciding with human colonisation events of the Seychelles archipelago. Contemporary genomes are over one-third inbred from recent inbreeding measured by runs of homozygosity (ROH) formed since human colonisation of the Seychelles, 50 generations ago (mean *F*_ROH < 50 generations ago_ = 0.38), and we identify significant inbreeding depression across cross key fitness traits. On average, a 17.9% reduction in lifespan and a 15.1% reduction in lifetime offspring production were associated with each 10% increase in individual inbreeding. Genome-wide scans reveal this depression is caused by a polygenic load. Our results suggest that a history of small population size could have facilitated the selective purging of severe genetic load, while mildly deleterious alleles escaped selection via drift. A partitioned genetic load architecture potentially enabled species recovery despite the inbreeding depression that followed near-extinction.

## Introduction

Populations under extreme bottlenecks risk entering an extinction vortex (Keller & Waller, 2002). In shrunken populations, increased inbreeding, the mating between relatives, exposes deleterious recessive alleles in homozygous state, and reduces fitness (Charlesworth & Willis, 2009). This inbreeding depression, coupled with reduced genetic diversity and vulnerability to stochastic events, is predicted to lead to population collapse (Hedrick & Kalinowski, 2000). Understanding the genetic and fitness consequences following a population bottleneck is a central question in conservation biology. This is difficult to quantify in populations that go extinct, as they are, by definition, unsamplable. Alternatively, quantifying the genomic architecture of inbreeding in surviving post-bottlenecked populations may yield evidence as to their persistence.

Whether a population survives a bottleneck despite intense inbreeding likely depends on the nature of the deleterious recessive alleles segregating in the population, otherwise known as its genetic load (Kardos et al., 2021). The genetic load of a population is often determined by its demographic history. Frequent inbreeding in small populations can eliminate or “purge” the most severely deleterious recessive alleles through selection, theoretically priming a population for survival through a crash (Hedrick & Garcia-Dorado, 2016). However, mildly deleterious recessive alleles are expected to escape natural selection and drift to high frequencies (Dussex et al., 2023). These mildly deleterious alleles that escape selection by drift during a bottleneck poses a long-term fitness cost to a population. Understanding the genomic architecture of inbreeding in populations that survive bottlenecks is a key goal which could enable prediction of conservation outcomes of other threatened species (Kyriazis et al., 2025). Resolving this requires a rare combination of data: an extreme, precisely dated bottleneck event, high-density genomic markers to map inbreeding, and accurate fitness monitoring with large sample sizes to power quantification of resulting effects.

The isolated population of Seychelles warblers (*Acrocephalus sechellensis*) on Cousin Island provides an excellent model-system to investigate the genomic architecture of inbreeding following an extreme bottleneck. The species underwent a catastrophic decline following human colonisation of the Seychelles archipelago in 1770, culminating in a mid-20th-century bottleneck of under 30 individuals (Vesey-Fitzgerald, 1940; Crook, 1960; Loustau-Lalanne, 1968). Critically, this population offers exceptionally accurate fitness quantification by virtue of its small and closed population that has been subject to detailed individual-based monitoring for 98% of all individuals since 1982 (Komdeur, 1992).

Here, we leverage a recently developed 1.8-million SNP genomic toolkit from 1,935 individuals sampled over 37 years to reconstruct the species’ demographic trajectory and characterize genomic inbreeding signatures of the bottleneck. By partitioning autozygosity into generation-specific classes, we evaluate the strength of inbreeding depression across four key fitness proxies: annual survival, lifespan, annual fecundity, and lifetime offspring produced. Finally, we conduct genome-wide scans to determine loci implicated in inbreeding depression and ongoing purging of recessive lethal haplotypes. Our research provides empirical evidence of the genomic architecture of inbreeding in a species surviving an extreme bottleneck.

## Methods

### Seychelles warbler population of Cousin Island

Seychelles warblers are facultative cooperative breeders (Hammers et al., 2019; Komdeur, 1991) with long lifespans (mean: 4.6 years from fledging, sd: 4.1, maximum: 20 years (Table S1)). They are endemic and culturally significant to the Seychelles. Between the 1940s and 1960s, the global population of Seychelles warblers was reduced to an anecdotal count of 26 (Crook, 1960; Vesey-Fitzgerald, 1940) within the 0.2-hectare mangroves of 29-hectare Cousin Island (04° 19.8′ S, 55° 39.8′ E). In 1968, BirdLife International purchased the Island, replacing coconut plantation with pan-tropical coastal vegetation, providing suitable habitat for the Seychelles warbler population to ultimately recover to 240–430 individuals (Komdeur, 1992, 1994; Komdeur & Pels, 2005). Translocations from Cousin have founded viable populations on four other islands (Richardson et al., 2006; Wright et al., 2014), resulting in a total, global population of >3,000 individuals (Brown et al., 2023).

The Cousin Island warbler population has been monitored for over three decades (Komdeur, 1992). Since 1997, >96% of all individuals have been ringed with a unique combination of a BTO metal ring and three color rings enabling identification of individuals (Komdeur et al., 1997; Raj Pant et al., 2020; Richardson et al., 2001). Individuals are usually first caught as nestlings, or as dependent juveniles (<5 months old) in their natal territory using mist nets (Kingma et al., 2016). Fieldwork takes place twice a year during the major (June to September) and minor (January to March) breeding seasons. The population is small and mostly contained, with only one Seychelles warbler from Cousin, having been recorded to migrate between islands in the archipelago (Komdeur et al., 2004). This enables high annual resighting rates for every individual. A blood sample, among other phenotypic measurements, is taken for each ringed individual, enabling accurate parentage assignment from genotype data (Lee et al., 2026). High annual resighting rates and parentage assignment enable exceptionally accurate estimation of fitness measures such as annual survival, lifespan, annual fecundity and lifetime fecundity.

### Genomic data

We used 1,929 Seychelles warblers from the Cousin Island population sequenced at low coverage (mean = 2.65×, SD = 1.70) using blood samples collected since 1982 and stored in absolute ethanol at −20°C to generate a raw genomic dataset (Figure S1). Reads were mapped to the chromosome-level reference genome, with accompanying functional annotations (Lee et al., 2026). We called 18 million variants and imputed them using STITCH, to at least 96.2% accuracy, as confirmed by downsampling to 0.1× coverage and comparing to a high-coverage reference set (Lee et al., 2026). We created a pedigree using 572 autosomal SNPs in sequoia v2.5.6 (Huisman, 2017), filtered for minor allele frequency > 0.3, genotyping rate > 99.9 and linkage (--indep-pairwise 1000 2 0.1 in PLINKv2.0; (Chang et al., 2015), and excluding SNPs on chromosomes with <90% imputation accuracy. We sexed samples using (1) proportion of called W chromosome genotypes (where females are expected to have a high value), calculated using PLINK v2.0’s function --missing, (2) Z chromosome coverage scaled to genome-wide coverage (where females are expected to have a lower value), calculated using the idxstats function in samtools v1.11 and dividing by mean sample coverage, and (3) Z chromosome heterozygosity (where females are expected to have values close to zero). Full details of the genomic data and pedigree are provided in Lee et. al (2026).

### Demographic history construction

To reconstruct the recent demographic history of Cousin Island Seychelles warblers, we sampled from the imputed genomic dataset 11 million autosomal SNPs, across 1,657 individuals, that passed filtering for a call rate > 0.999 and individual genotyping rate > 0.96 in *GONE* (Santiago et al., 2020). *GONE* estimates effective population size (*N*_e_) by calculating LD between pairs of SNPs, and can estimate demographic history up to 100 generations in the past, using a maximum of 1,800 individual samples. The closed-island population of Seychelles warblers on Cousin is well suited to this analysis, which assumes no admixture in the population. Specific to *GONE*, we used the following parameters: 1 cM/Mbp recombination rate, 2,000 generations, 400 bins, a maximum of 20,000 SNPs per chromosome, no minor allele frequency (MAF) pruning and allowing Haldane correction. We ran 50 iterations each with a new subsample of 20,000 SNPs (Shaw et al., 2025) and then plotted the mean *N*_e_ across runs (Fig. 1a). Reported Seychelles warbler generations were converted to years by calculating the average age of parents in our dataset, giving a generation time of 4.8 years.

**Figure 1:**
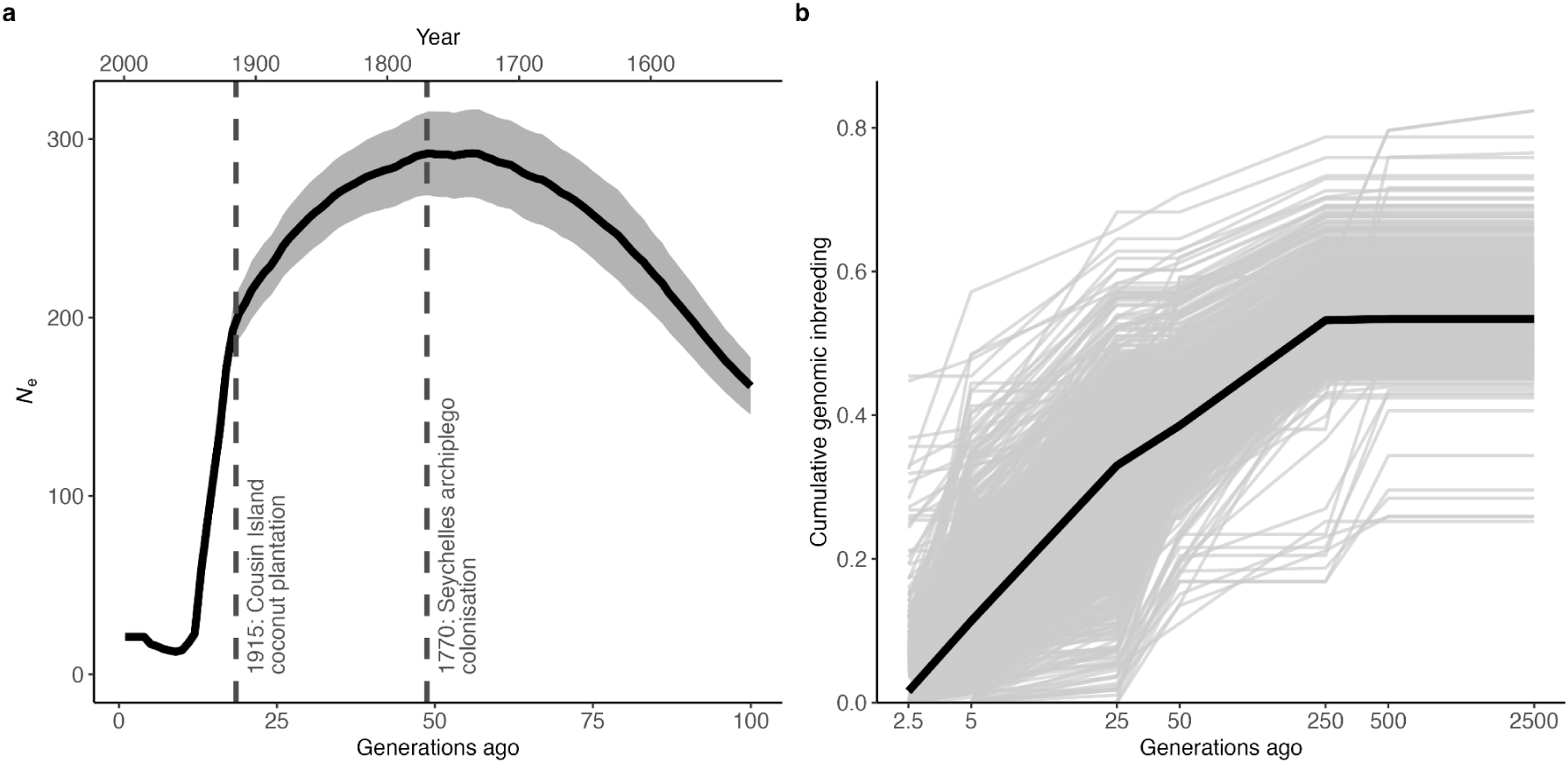
Seychelles warblers demographic and inbreeding history on Cousin Island, Seychelles. **a**. Estimated mean effective population size (solid line) with 95% confidence intervals (shaded area) over the past 100 generations (generation time = 4.8 years) based on LD between whole-genome SNP data from 1,657 Seychelles warbler from Cousin island. Key historical events are denoted by overlaid vertical dashed lines. **b**. Accumulation of genomic inbreeding, ROH (y-axis) binned by age of ROH (x-axis, log scale) per individual (1,929 grey lines) and for the mean of all individuals (orange line).

### Inbreeding coefficients

We used 1,890,803 autosomal SNPs, filtered for call rate > 0.99 and MAF >0.01 across 1,929 individuals, that passed filtering for mean coverage > 0.05, yielding a genome-wide SNP density of 1,870 SNPs/Mbp and sample-wide genotyping rate of 0.998. We quantified individual inbreeding coefficients using the hidden Markov model-based approach to identify ROH segments (Druet & Gautier, 2017) in the R (R Core Team, 2022) package RzooRoH v.0.3.2.1 (Bertrand et al., 2019). This model-based approach incorporates genotype probabilities of our imputed SNPs when quantifying individual inbreeding (partitioned by ROH segment length). It is robust to genotyping density and error rate when compared with rule-based approaches that require user-defined parameters to qualify a ROH (Shafer & Kardos, 2025). To model individual inbreeding, we used nine ROH classes that corresponded to the number of generations since the most common recent ancestor of the ROH segment (2.5, 5, 25, 50, 250, 500, 2,500 and 7,500 generations ago) and one non-ROH class (Figure S1), using a genotyping error rate of 1% (Lee et. al 2025). The model qualifies regions as being ROH when the ROH probability is > 0.99. We plotted the ancestral accumulation of inbreeding coefficients from each ROH class (Fig. 1b). We also plotted the mean density distribution of all ROH segments in 500-kbp windows across the genome (Fig. S2).

To quantify inbreeding depression, we estimated individual inbreeding coefficients (*F*_ROH_) as the sum of the first four ROH classes above, divided by autosomal genome size, as these captured “recent inbreeding” since the 1770 European colonisation of Seychelles, up to ∼50 generations ago, assuming the generation time for Seychelles warblers is 4.8 years (Spurgin et al., 2014). We plotted the *F*_ROH_ distribution among individuals (Fig. S3a), across annual cohorts (Fig. S3b) and of those alive as of 2025 (Fig S4). To evaluate robustness, we verified the correlation of individual *F*_ROH_ estimates with those generated by a rule-based approach in PLINK v.2.0 (Fig. S5a), and with (Fig. S5b). In PLINK v.2.0, we used the following parameters: --homozyg-window-snp 50 --homozyg-snp 50 --homozyg-kb 1000 --homozyg-gap 300 --homozyg-density 200 --homozyg-window-missing 4 --homozyg-het 2 --homozyg-window-het 2. To ensure comparability with the recent *F*_ROH_ estimates from our HMM-based approach, we specified a minimum ROH length of 1 Mbp. This corresponds to inbreeding originating approximately 50 generations ago, calculated as *L* = 100 / (2*g*), where *g* = 50 generations and assuming a genome-wide recombination rate of 1 cM/Mbp (Thompson, 2013). We estimated pedigree-based inbreeding coefficients in the R package FnR v.1.1.0 (Nilforooshan, 2024) for individuals that had both their parents and maternal grandparents known (*n* = 1,104).

### Inbreeding depression models

We investigated inbreeding depression in Seychelles warblers by testing covariation between *F*_ROH_ and four single-generation fitness proxies (Table 1): annual survival, lifespan, annual fecundity and lifetime fecundity. These metrics correlate strongly with multi-generational fitness in wild populations (Brommer et al., 2004; Hunter et al., 2019; Reid et al., 2019; Van de Walle et al., 2022; Young et al., 2023), including in the Seychelles warbler (Chesterton et al., 2024). We fitted generalized linear mixed models (GLMMs) using a Bayesian framework implemented in the R package MCMCglmm (Hadfield, 2010). We restricted the analysis of lifetime fitness traits to individuals with completed life histories by excluding 72 birds still alive in the dataset after 2024, and excluding 115 translocated birds, leaving *n* = 1,735 individuals for the annual models and *n* = 1,663 for the lifetime models. We ran separate models for each fitness trait:

1. Annual Survival: Modeled as a binary variable (0 = died, 1 = survived to next year) using a “threshold” family.
2. Annual Fecundity: Modelled as the number of genetically verified offspring (including extra-pair) produced in a given year, using a Poisson distribution.
3. Lifespan: Modelled as the age at death, using a Poisson distribution.
4. Lifetime fecundity: Modelled as the total number of genetically verified offspring produced over an individual’s life, using a Poisson distribution. We distinguish this metric from Lifetime Reproductive Success (LRS), which typically requires offspring recruitment into the breeding population.

For all Poisson models, we included an observation-level random effect to account for overdispersion (Chesterton et al., 2024; Sparks et al., 2022). The primary predictor of interest was individual inbreeding, *F*_ROH_. To facilitate biological interpretation, we rescaled *F*_ROH_ from 0–1 to 0–10. Consequently, model estimates reflect the change in fitness resulting from a 10% increase in genome-wide homozygosity (Stoffel et al., 2021). To test for environmental modulation of inbreeding depression, we included an interaction term between *F*_ROH_ and environmental harshness. For annual models, harshness was defined as the annual mortality rate (total deaths / total population size) in the year of observation. For lifetime models, we used the mortality rate in the individual’s birth year.

We controlled for confounding variables identified in previous Seychelles warbler studies: sex (Richardson et al., 2004; Sparks et al., 2022), standardised maternal age (Richardson et al., 2004; Sparks et al., 2022) and age, which in annual models, we included both linear and quadratic (Age^2^) terms to account for senescence (Brown et al., 2022; Hammers et al., 2015; Komdeur, 1996). All models included an additive genetic random effect linked to the pedigree to control for heritability of fitness traits and to account for the non-independence of observations due to relatedness. Annual models included individual identity to account for repeated measures across an individual’s life, and birth year to control for cohort effects. To prevent the overestimation of effect sizes and artificial narrowing of credible intervals associated with stepwise variable selection, we retained all specified fixed and interaction terms within our final models regardless of statistical significance, interpreting the effect sizes of *F*_ROH_ solely from these full global structures (Forstmeier & Schielzeth, 2011; Whittingham et al., 2006). We verified model convergence by visually inspecting trace plots and observing low autocorrelation among posterior samples and confirming that all core parameters passed the Heidelberger and Welch stationarity diagnostic in the coda R package. We translated model estimates into biological effect sizes corresponding to a 0.1 increase in *F*_ROH_ (a 10% increase in genome-wide homozygosity). For traits modelled with a Poisson distribution (log-link), we calculated the percentage change in the fitness metric as (e^β^ - 1) x 100. For annual survival (probit-link), we calculated the absolute change in survival probability relative to the mean adult (>1 years old) survival rate observed in the population. To contextualise our findings, we converted our effect size and those from comparable bird studies into correlation coefficients (*r*) and Fisher’s *z* scores following standard protocols (Nakagawa & Cuthill, 2007).

### ROH genome-wide association scan (GWAS)

We tested, for each allele of a SNP in a ROH, its association with lifespan and lifetime fecundity (Duntsch et al., 2023; Stoffel et al., 2021). For each of the 1.8 million SNPs, we tested covariance between survival and the effect of being homozygous for each allele within a ROH, and extracted their estimated slopes and *p*-values using lme4 (Bates et al., 2015). We controlled, as fixed effects, the additive effects of the SNPs, regardless of whether they were in a ROH, background *F*_ROH_ for the non-focal chromosomes, overall genetic background as measured by the top five principal components of the variance-standardised additive relationship matrix (PC1–5), sex, death rate of the birth year and standardised maternal age. We assumed negative binomial error distributions. To determine threshold significance, we estimated the effective number of independent SNPs with a linkage disequilibrium value of r^2^ < 0.8 between SNPs in 5,000-kbp regions as 10,994 effective SNPs. As we conducted two tests per model we doubled this number to 21,988 and applied Bonferroni correction of *p*-values, resulting in a threshold significance of *p* < 2.27 * 10^-6^. We identified genes implicated in high-impact SNP regions using their overlap in position with the Seychelles warbler functional annotations (Lee et al., 2026).

### Recessive lethal haplotypes

We conducted a genome-wide scan for recessive lethal haplotypes to investigate whether early-life mortality occurring prior to genetic sampling has potentially purged individuals carrying high-effect deleterious mutations. This is because natural nesting sites of the Seychelles warbler are often located high in trees and so preclude the collection of comprehensive clutch-level data.

We first phased the imputed genotype data to reconstruct chromosomal haplotypes. We phased data using SHAPEIT5 (v5.1.1) (Hofmeister et al., 2023), which utilises both population-level linkage disequilibrium (LD) and pedigree information to resolve phase. Given the lack of a species-specific recombination map, we used a constant recombination rate based on a conservative effective population size (*N_e_*) of 1,000, reflecting the species’ recent bottleneck history. The use of pedigree information (1,539 trios), with a low rate of Mendelian inconsistencies of < 0.03%, allowed for phasing. To capture Identity by Descent (IBD) segments while maximising genomic resolution, we defined haplotypes using sliding windows of 1000 SNPs, corresponding to a physical distance of approximately 550 kbp. Windows were slid one SNP at a time across all autosomes for fine-scale mapping of potential lethal loci.

For each defined haplotype window, we identified all unique haplotypes with a population frequency > 0.5%. We then tested for a deficit of homozygous offspring (Stoffel et al., 2023). For each unique haplotype (*h*), we identified all matings in the pedigree where both parents were carriers (genotype *Hh*). We calculated the expected number of homozygous offspring (*E*_hh_) based on Mendelian transmission probabilities, assuming independent assortment.

A chi-square test was used to compare the observed number of homozygous offspring (*O*_hh_) to the expected number (*E*_hh_). To ensure statistical robustness, we only tested haplotypes where *E*_hh_ ≥ 5. The test statistic was calculated as:

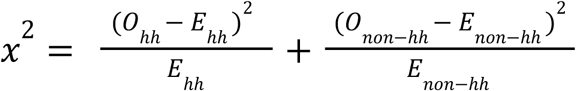

where non-hh represents all other genotype classes (*HH* and *Hh*). We specifically tested for a deficit of homozygotes by filtering for results where *O*_hh_ < *E*_hh_. Initial genome scans revealed large, highly significant peaks on chromosome 3 that were coincident with regions of anomalous sequencing coverage, and corresponded to the same region where ROH could not be called to (Fig. S2), suggesting a local reference genome misassembly. To prevent these technical artifacts from confounding the analysis, we excluded any haplotype window overlapping these regions.

## Results

### Demographic history construction

Linkage disequilibrium (LD) analysis of 11 million imputed autosomal SNPs from 1,657 individual Seychelles warblers on Cousin Island using *GONE* suggested a decline in effective population size, *N*_e_, from ∼200–300 starting ∼50 generations ago (year ∼1780), reaching a minimum *N*_e_ of ∼13 only ∼9 generations ago (year ∼1960) that has since remained low (Figure 1a).

### Inbreeding

Quantification of inbreeding by hidden Markov model (HMM) from 1.8 million imputed SNPs across 1,929 individuals identified a total of 1,082,777 homozygous-by-descent (HBD) segments (mean length: 0.92 Mbp; maximum length 154 Mbp), which we refer to hereafter as ROH for sake of consistency with previous literature. ROH segment density in 500-kbp windows varied considerably across the genome, and almost entirely absent in a missassembled 9-Mbp region of Chromosome 3 and in Chromosome 19 (Fig. S2). A majority, 72.8% of ROH from the average Seychelles warbler genome have accumulated over the most recent ∼50 generations, which is just after European colonisation of Seychelles in ∼1770 (Fig. 1a). Summing all 182,132 ROH classes that captured recent inbreeding up to ∼50 generations ago, and dividing by autosomal genome size, gives a mean *F*_ROH_ estimate of 0.38 per individual (SD: 0.057, Fig 1b). *F*_ROH_ did not change over our 37-year dataset (Fig. S3b). *F*_ROH_ was strongly correlated with the PLINK-derived rule-based *F*_ROH_ (*r* = 0.93, Fig. S5a) and less so with pedigree-based inbreeding (*r* = 0.55, Fig. S5b). We display the *F*_ROH_ distributions by ROH segment length for 65 whole-genome sequenced individuals that were still alive in 2025 as examples of potential candidates for future translocations (Fig. S6).

### Inbreeding depression

Seychelles warblers showed significant evidence of inbreeding depression, as estimated by whole-genome estimation of *F*_ROH_, in all four fitness components tested (Table 1): annual survival (Fig. 2a), lifespan (Fig. 2b), annual fecundity and lifetime fecundity (Fig. 2c). Full model summaries from the Bayesian regression analyses that control for confounding variables are provided in Table S2.

**Figure 2.**
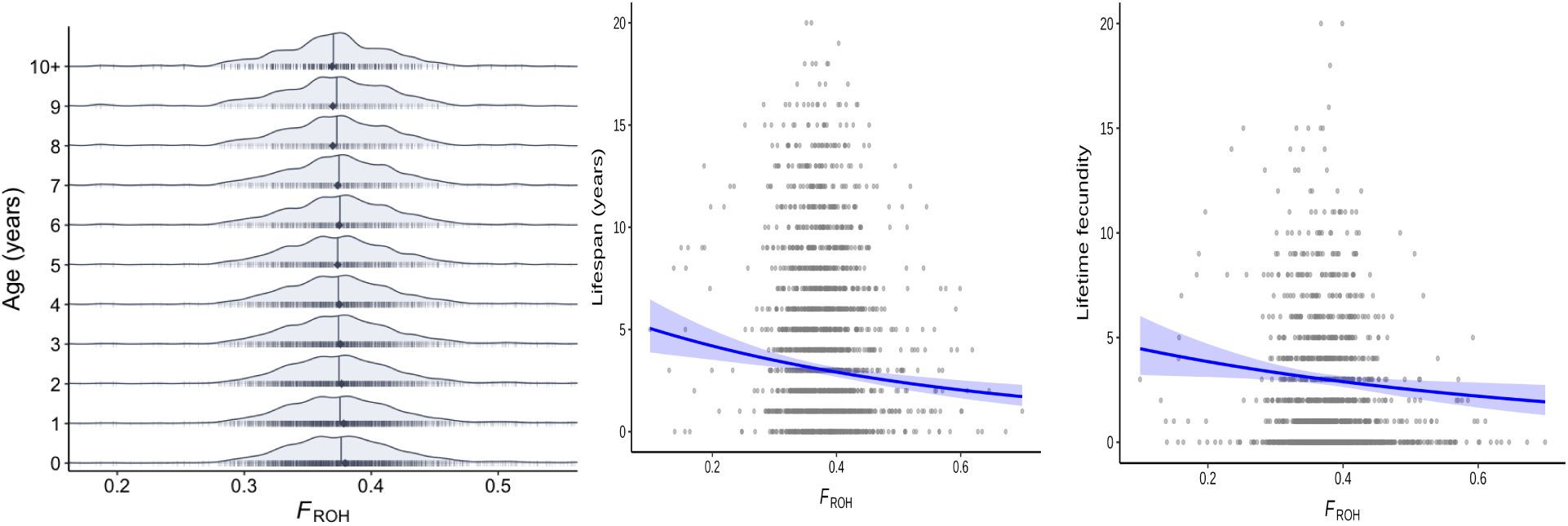
Fitness effects of inbreeding on survival and reproduction. **a.** *F*_ROH_ distributions among age classes ranging from 0 to 10+ years, showing median (vertical lines) and mean (diamond points) values. **b.** *F*_ROH_ distributions by lifespan. **c.** *F*_ROH_ distributions by lifetime fecundity. The shaded blue provides 95% credible intervals for predicting fitness by *F*_ROH_ while holding the following effects constant: mean birth year, death rate and mean maternal age.

Annual island mortality rate covaried significantly with annual fitness, but birth year mortality rate did not covary with lifespan or lifetime fecundity. These mortality rate measures, which we included as proposed proxies for island environmental quality, did not significantly influence the inbreeding depression relationship in any model. Linear increases in age were associated with reduced annual survival, and also with increased annual fecundity, though the quadratic age term suggests that annual fecundity decreased in later life. Individuals with older mothers showed significantly reduced annual survival and lifespan but not annual fecundity or lifetime fecundity. We only found a sex difference in annual fecundity. Additive genetic components to fitness were consistently low, and even lower for fecundity fitness traits.

**Table 1.**
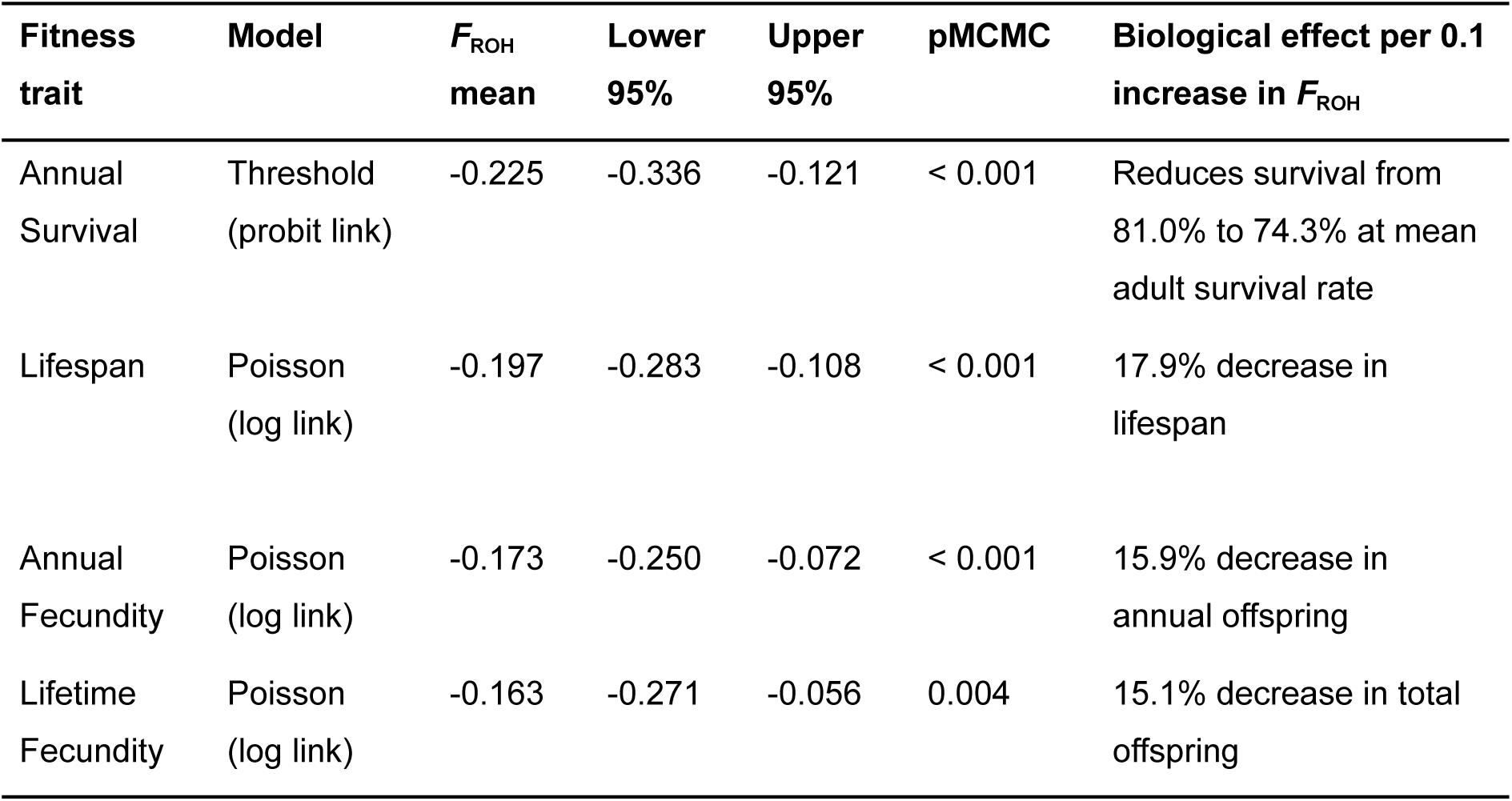
Summary of inbreeding (*F*_ROH_) effects on four fitness components in Seychelles warblers from Bayesian regression analysis. Shown are the fitness trait, error distribution, scaled posterior mean of F_ROH_, 95% credible intervals, a Bayesian analogue of the frequentist *p*-value (pMCMC), and the biological interpretation a 0.1 increase in *F*_ROH_. Values are extracted from each of the four fitness models in Table S2, which details outputs for covariates.

| **Fitness trait** | **Model** | **$F_{ROH}$ mean** | **Lower 95%** | **Upper 95%** | **pMCMC** | **Biological effect per 0.1 increase in $F_{ROH}$** |
| --- | --- | --- | --- | --- | --- | --- |
| Annual Survival | Threshold (probit link) | -0.225 | -0.336 | -0.121 | < 0.001 | Reduces survival from 81.0% to 74.3% at mean adult survival rate |
| Lifespan | Poisson (log link) | -0.197 | -0.283 | -0.108 | < 0.001 | 17.9% decrease in lifespan |
| Annual Fecundity | Poisson (log link) | -0.173 | -0.250 | -0.072 | < 0.001 | 15.9% decrease in annual offspring |
| Lifetime Fecundity | Poisson (log link) | -0.163 | -0.271 | -0.056 | 0.004 | 15.1% decrease in total offspring |

### Genome-wide association with inbreeding effects

Our mixed models tested the fitness effect of each allele in a SNP being homozygous within a ROH for lifetime fecundity (Fig. 3) and lifespan (Fig. S6). Distribution plots of effect sizes (Fig. 3a, Fig. S6a) and *p*-values (Fig. 3b, Fig. S6b) suggest that most homozygous SNPs within ROH are deleterious and when deleterious, showed larger effect sizes than when beneficial. We did not detect any SNPs that passed the genome-wide significance threshold, though some approached the significance threshold. This included a signal impacting lifespan on chromosome 3 between positions 43,295,985–43,300,864 bp that overlapped with the *AGPAT4* gene, an enzyme involved in phospholipid synthesis. We also found two near-significant signals reducing lifetime fecundity. On chromosome 2, a region between positions 3,178,652–3,199,945 bp overlapped with the *ADAMTS1* gene, a metalloproteinase that breaks down the extracellular matrix. On chromosome 9, a region between positions 10,937,079–11,556,800 bp overlapped with the *SLC6A6* gene, a protein that transports taurine.

**Figure 3.**
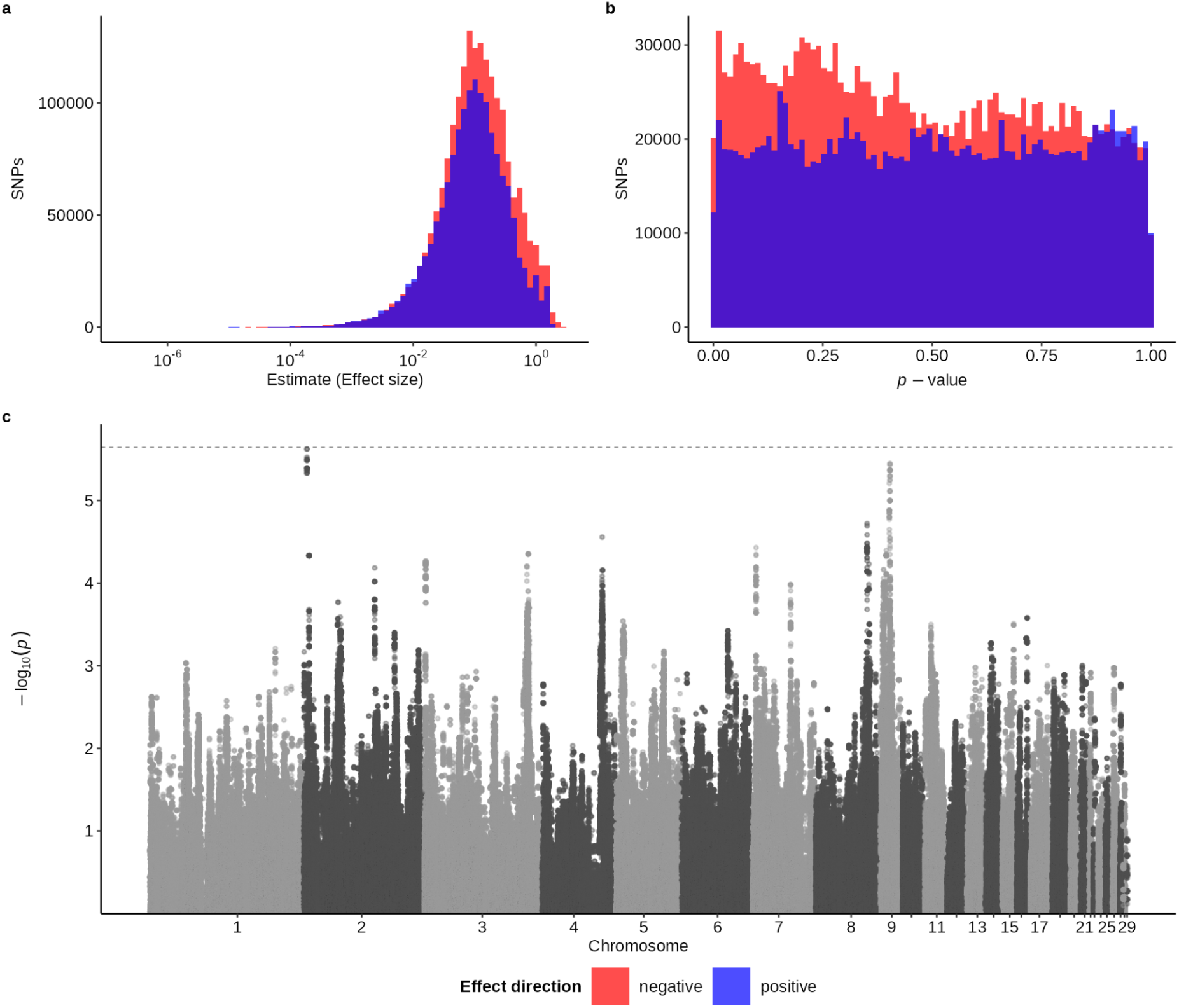
GWAS of ROH status effects on lifetime fecundity. **a.** Distribution of effect sizes for each SNP’s allele within ROH. **b.** Distribution of *p*-values for SNPs within ROH. **c.** Manhattan plot of ROH status *p*-values across the genome. The dashed line represents the genome-wide significance threshold.

### Recessive lethal haplotypes

Despite 1,929 samples, a deep pedigree (1,539 trios), and dense genotyping (1,890,803 SNPs and 18,452 unique haplotypes), we found no evidence of recessive lethal or semi-lethal haplotypes. The genome-wide scan for homozygote deficiency showed that all haplotypes fell below the significance threshold (Fig. 4). The haplotype candidate closest to significance was on Chromosome 13 (window start: 10,414,842), where we observed zero observed homozygotes, despite an expected 16.25 based on a carrier count of 313 (𝜒^2^ = 21.90, *p =* 2.87 * 10^-6^).

**Figure 4.**
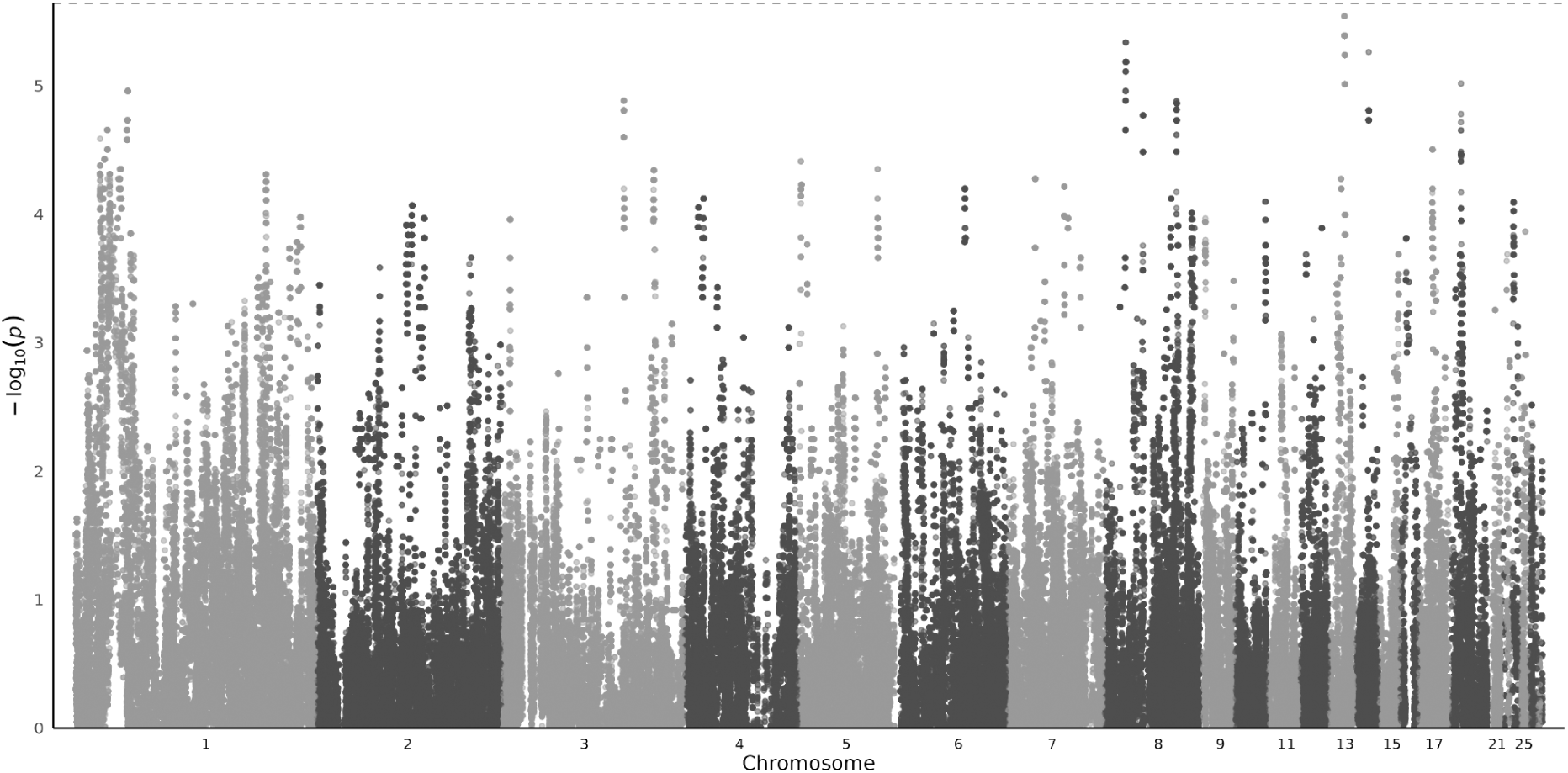
Genome-wide sliding-window scan for recessive lethal haplotypes. The y-axis represents *p*-values derived from chi-square tests comparing observed and expected homozygote counts for unique haplotypes (1,000 SNP windows ∼ 550 kbp, carrier frequency > 0.5%). The dashed line represents the genome-wide significance threshold.

## Discussion

Seychelles warblers have recently been bottlenecked in the last surviving population on Cousin Island, Seychelles. The inbreeding that followed this population decline measured by whole-genome estimates of *F*_ROH_, resulted in inbreeding depression on four key individual fitness traits, measured from long-term monitoring: annual survival, lifespan, annual fecundity and lifetime fecundity. Our results suggest that inbreeding depression has arisen due the negative effects of many weakly deleterious alleles being expressed when in the homozygous form within ROH, with little evidence for ongoing purging of lethal recessive haplotypes.

There is an overall pattern of a very recent population crash starting 50 generations ago, in ∼1780, that closely follows key historical events following the colonisation of Seychelles. In 1770, European settlers claimed Seychelles for its resources, including timber and the cultivation of spices, crops, cotton and, later, coconut, while introducing many non-native wild species (Allen, 2022; Monnier, 2019; Swabey, 1970). Indigenous forest was intentionally burned by traders to control the Coco-de-Mer market (Etongo et al., 2021). By around 1915, Cousin island, adjacent to Praslin, became a coconut plantation (Komdeur & Pels, 2005; Scarr, 2000; von Brandis, 2012) and ultimately held the remaining population of individual Seychelles warblers that had survived on Cousin Island with an *N*_e_ as low as ∼13 from a pre-Anthropogenic *N*_e_ of ∼250–300. Although recent demographic history inference is possible, as noted by the authors of the analysis package, *GONE, N*_e_ trends in the more distant past should be treated as less exact because of the impact of genetic recombination eroding linkage disequilibrium patterns between SNPs (Santiago et al., 2020). These results corroborate an earlier study that compared *N*_e_ in museum samples (mean collection year = 1898) and contemporary Seychelles warblers (Spurgin et al., 2014) using 12 microsatellite markers. Our *N*_e_ estimates align more closely with the ONeSAMP estimates (museum population: mean = 268, 95% credible intervals = 175–1320; contemporary population: mean = 32; 95% credible intervals = 27–41) than their Approximate Bayesian Computation (ABC)-derived ancestral estimates (museum population: median = 6,900, 95% credible intervals = 2,400–9,700; contemporary population: median = 46; 95% credible intervals = 29–75). It is possible that the larger *N*_e_ estimated by ABC may be heavily influenced by the broad, prior assumptions regarding ancestral population sizes, which were defined by Spurgin *et al*. (2014) as 1–100,000.

Accumulation of inbred ROH segments in the average Seychelles warbler genome closely follows expectations based on the genomic analysis of demographic history. A majority of 72.8% of ROH from the average Seychelles warbler genome have accumulated over the most recent ∼50 generations. We thus investigated inbreeding that occurred up to ∼50 generations ago (∼year 1770), the beginning of human colonisation in the Seychelles, to define *F*_ROH_. Distinguishing ROH segments originating specifically from the last 50 generations allowed us to isolate the genetic consequences of inbreeding that occurred since the human-mediated bottleneck. The hidden Markov model based estimate of *F*_ROH_ we used correlated well with rule-based-estimation of *F*_ROH_ (*r* = 0.93) and reasonably well with inbreeding estimates generated by the SNP-assembled pedigree (*r* = 0.55). These correlations were similar to those reported in other studies comparing model-based and rule-based genomic inbreeding estimates (Duntsch et al., 2023; Lavanchy & Goudet, 2023) and genomic-based and pedigree-based inbreeding estimates(Bem et al., 2024; Cortes-Hernández et al., 2021; Huisman et al., 2016). It is advantageous to use a model-based approach (RZooRoH) to call ROH, as we can probabilistically partition autozygosity into distinct age classes, providing a more robust estimation of recent inbreeding that is less sensitive to genotyping error and varying marker density than rule-based methods (Wæge et al., 2025).

Over a third (*F*_ROH_ = 0.38) of the average Seychelles warbler genome constitutes ROH formed since human colonisation at around 1770, or 50 generations ago. The density of these ROH regions varies across the genome. Dense ROH regions in the genome may coincide with recombination “coldspots”, or harbour loci that have undergone recent hard sweeps under positive selection. Conversely, regions of low ROH density may have previously harboured strongly deleterious alleles that would have been purged in the homozygous state, becoming systematically absent from sequenced survivors. We noted a complete absence of ROH density across samples in a ∼9-Mbp region spanning 15.1 –23.8 Mb on chromosome 3 and chromosome 19. The region on chromosome 3 coincided with anomalously high sequencing coverage, indicating a local genome misassembly, such as a collapsed repeat. In contrast, the lack of ROH on chromosome 19 is attributed to its poor assembly quality, stemming from the known evolutionary dynamism of this chromosome within the *Sylvioidea* (Lee et al., 2026). Although these regions were not excluded from the global inbreeding depression models, their impact on individual *F*_ROH_ estimates will be negligible, as they represent less than 1% of the total autosomal assembly, and will be consistent across all individuals.

The sampled Seychelles warbler population showed significant evidence of inbreeding depression in all four tested single-generation proxies of fitness: annual survival, lifespan, annual fecundity and lifetime fecundity. Based on our model estimates, a 0.1 increase in *F*_ROH_ (representing a 10% rise in genome-wide homozygosity) reduced lifespan by 17.9% and total lifetime offspring by 15.1%. Despite potentially expecting compounding effects on annual survival and annual fecundity, the effect size is slightly stronger for lifespan, likely due to the environmental stochasticity inherent in lifetime fecundity such as mate quality, which dilutes the genetic signal relative to the total phenotypic variance. Overall inbreeding costs are similarly expressed in both survival and reproductive traits.

These effect sizes are comparable with other long-term monitored bird systems that have modelled covariation between among other fitness traits, lifetime fecundity and genome-based inbreeding coefficients (Table 3). We note that other studies often have access to clutch-level data via nest-box monitoring, whereas Seychelles warblers use high, natural nests that are mostly inaccessible. Consequently, our estimates mainly quantify fitness in fledged individuals.

**Table 2.**
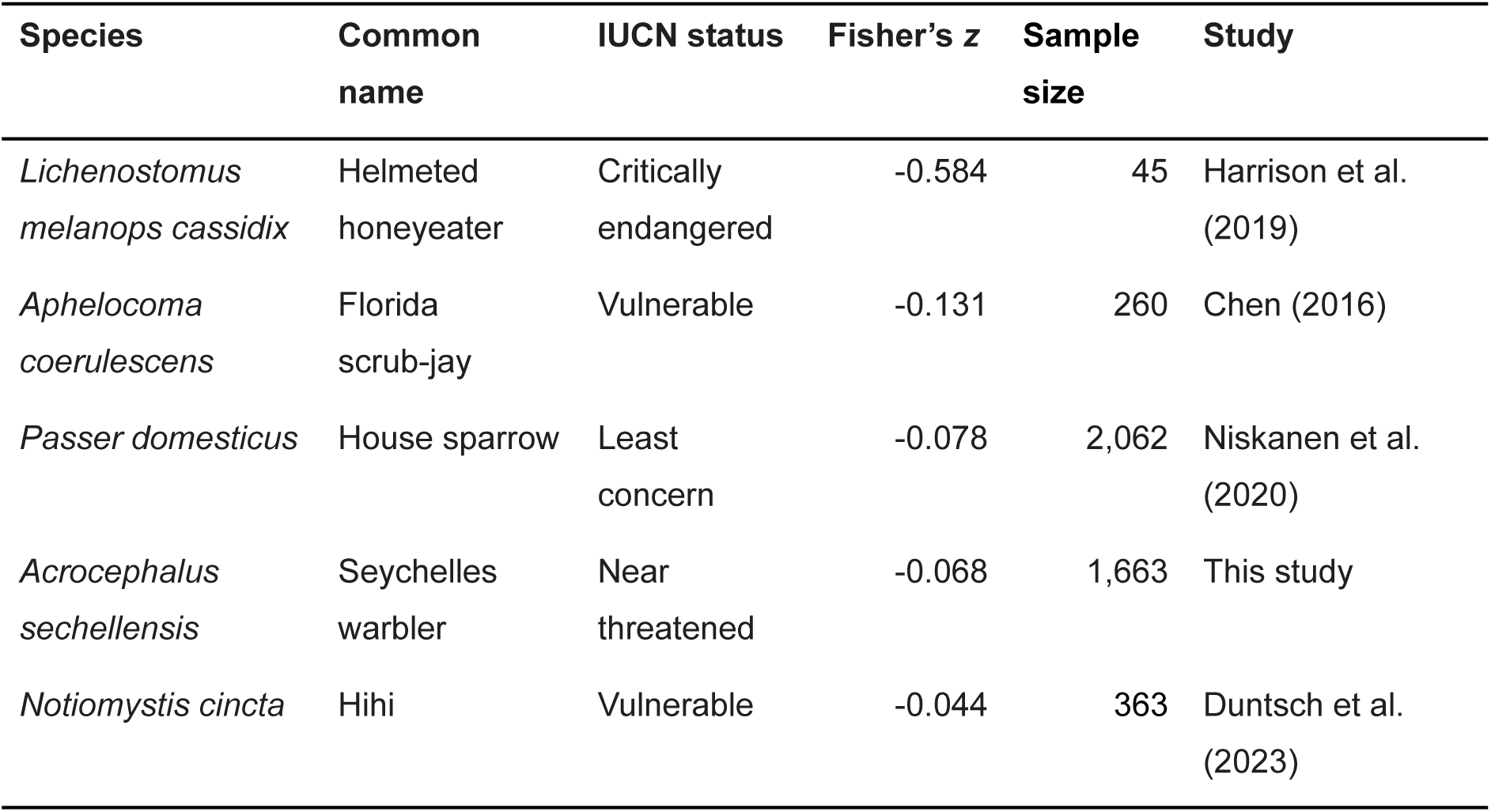
Comparison of standardised inbreeding depression effect sizes across long-term monitored bird populations. Effect sizes (Fisher’s *z*) represent the correlation between genome-wide inbreeding and lifetime offspring production. A more negative z value indicates a stronger deleterious effect of inbreeding on fitness.

Importantly our findings contrast with previous investigations in this system using heterozygosity-fitness analyses that were typical of the time, but likely lack the resolution of genomic inbreeding estimates (Balloux et al., 2004; Slate et al., 2004). Earlier work using 14 microsatellite markers found no effect of heterozygosity on survival (*n* = 119, Richardson, Komdeur and Burke, 2004), although a reduction in individual telomere length was later detected (*n* = 568, Bebbington *et al*., 2016). Our detection of significant inbreeding depression across all fitness traits is likely due to the improved power of our study, which used a near error-free SNP-based pedigree with 1,929 individuals and 1.8 million SNPs (Lee et al., 2026). The pervasive inbreeding depression identified here suggests strong selection for inbreeding avoidance, commonly linked to sex-biased dispersal in birds as an adaptation to avoid the costs of inbreeding (Pusey, 1987; Riehl & Stern, 2015). In Seychelles warblers, 40% of offspring result from extra-pair paternity, usually with nearby males, and so females may disperse further specifically to avoid inbreeding with local kin (Eikenaar et al., 2008). Future work could test this hypothesis with the prediction that extra-pair paternity offspring would have lower *F*_ROH_ than their within-pair half-siblings.

We found “harsher” years, as defined by a higher population-wide mortality rate, did not significantly exacerbate the deleterious effects of inbreeding on annual survival, annual fecundity or lifetime fecundity. The interaction for lifespan approached significance, but the overall trend suggests environmental stability regarding inbreeding costs. Importantly, our environmental proxy was biologically relevant: harsher years significantly reduced fitness across the population, where a 10% increase in annual mortality reduced life expectancy by ∼6% and annual fecundity by ∼9%. This absence of an interaction contrasts with earlier Seychelles warbler work, which reported that maternal inbreeding depression was mediated by breeding season quality (Richardson et al., 2004). That discrepancy likely reflects our improved resolution: we used high-density genomic data (*F*_ROH_), a sample size nearly twenty times larger (*n* = 1,663) and a continuous environmental proxy derived from 37 years of monitoring rather than a categorical season bin using only four breeding seasons. Our results align with a broader review of wild populations (Pemberton et al., 2017), where only 12 of 96 tests for inbreeding-by-environment interactions were significant, of which one test was the aforementioned Seychelles warbler study. It remains possible that interactions exist but are too subtle to detect, or that “harsh” years remove highly inbred individuals so that the interactive effect is not sampled.

As expected for fitness traits under strong selection (Postma, 2014), the additive genetic variance was small. By explicitly modeling this additive genetic component, we accounted for the non-independence of observations due to relatedness, ensuring that our estimates of inbreeding depression were isolated from background family effects (Lavanchy, Weir, et al., 2024; Yengo et al., 2017). Seychelles warblers had reduced survival and fecundity as they aged, though the fecundity effect appeared quadratic, with individuals initially increasing their fecundity when they were young (Brown et al., 2022). While a negative quadratic age term is highly suggestive of reproductive senescence, a formal post-peak or piece-wise regression analysis would be required to definitively confirm a late-life decline, an investigation that remains beyond the intended scope of this study. We confirmed the previously-found Lansing effects in Seychelles warblers (Sparks et al., 2022), that annual survival and lifespan reduced as maternal age increased, but there was no similar effect on annual fecundity or lifetime fecundity.

Our distributions of ROH effects show that any inbred region is more likely to have a negative than positive effect on lifespan and lifetime fecundity, and that negative effects have larger effect sizes and lower *p-*values. This suggests that in recently bottlenecked Seychelles warblers from Cousin island, inbreeding depression is highly polygenic, with many weakly deleterious alleles spread throughout the genome. Our search for recessive lethal haplotypes to investigate potential early-life mortality from inbreeding, that we could not sample directly in the nest, yielded no significant results from 1,539 pedigree trios. The strongest signal of a recessive lethal haplotype was found on chromosome 13, where 16 homozygous offspring were expected, and none were found, although this did not pass the significance threshold (Figure 4). Under extended periods of small *N*_e_, segregating lethal alleles are more frequently exposed to natural selection from inbreeding. Theoretical models suggest that populations with a long-term *N*_e_ < 500 are more efficient at purging highly deleterious mutations over evolutionary timescales (Kyriazis et al., 2021). Accordingly, our demographic reconstruction shows that *N*_e_ in the Seychelles warbler never exceeded 300, prior to human arrival, which may have primed the population for recovery by purging its most lethal recessive alleles and may explain the success of four subsequent translocations from Cousin Island to Aride (1988), Cousine (1990), Denis (2004), and Fregate (2011). The remaining inbreeding depression is polygenic, caused by the unmasking of many mildly recessive deleterious alleles across the genome that may have drifted to high frequencies, obscured by selection (Dussex et al., 2023; Kardos et al., 2021; Lavanchy, Cumer, et al., 2024). The patterns in the Seychelles warbler contrast with those reported in the St Kilda population of Soay sheep (Stoffel et al., 2021, 2023) and in the Tiritiri Mātangi island population of hihis (Duntsch et al., 2023), the only two other wild populations in which regional inbreeding depression at the whole-genome level has been quantified. Those studies show some evidence of harbouring significantly deleterious SNPs within ROH regions. Strongly deleterious SNPs may not yet have been purged in Soay sheep because of a recent admixture event (Feulner et al., 2013). In hihis, the Tiritiri Mātangi island population descends entirely from a very recent re-introduction in 1995, from their source population. The population potentially still retains highly deleterious recessive load brought from founding individuals (Brekke et al., 2011) and may explain high rates of reproductive failure. In hihi, a third of eggs fail to hatch (Morland et al., 2024) and supplementary feeding is currently suggested to be necessary for population persistence (Doerr et al., 2017; Roper & Brunton, 2024). This contrasts the 8% (13/157) hatching failure rates of Seychelles warblers (Komdeur et al., 2002).

## Conclusions

Our results demonstrate that after recovering from a recent bottleneck where *N_e_* fell to approximately 13, the Seychelles warbler continues to harbour a significant cost of inbreeding. By pairing 37 years of longitudinal field data with a high-density genomic toolkit, we identified pervasive inbreeding depression across four major fitness proxies, with a 10% increase in *F*_ROH_ reducing lifetime fecundity by 15.1%. The species’ long-term demographic history as an island endemic, with an effective population size that did not exceed 300 before human settlement may have facilitated a historical purging. Consequently, the contemporary genetic load in this system appears to be characterised by a polygenic accumulation of weakly deleterious alleles that are more resistant to natural selection, rather than a load of severely deleterious recessive alleles. To validate this, a temporal comparison of genetic load using 26 historical specimens held in museum collections is a logical next step. We would expect these museum samples to harbour a higher frequency of lethal recessive alleles that have since been purged in the contemporary population. Conversely, any weakly deleterious load may have increased due to genetic drift during the recent population crash. Finally, replicating the search for recessive lethals using an expanded pedigree is necessary to definitively confirm the absence of homozygous offspring on chromosome 13, where 16 were expected.

If the most lethal recessive haplotypes have been purged from Cousin Island, it may explain the population recovery of Seychelles warblers from the brink of extinction and subsequent successful establishment of new populations on Aride, Denis, Cousine, and Fregate islands by translocation. Any potential efforts to establish a fifth population could be optimised by prioritising translocation candidates with lower *F*_ROH_ to boost survival and reproductive prospects, as there is evidence of a remaining weakly deleterious load. Broadly, genomics has enabled quantifying a species’ demographic history to provide essential context to the underlying genetic load architecture. Studies of this nature are critical for predicting inbreeding costs in threatened wildlife, providing a high-resolution genomic blueprint that contextualizes genetic load dynamics in populations where such fine-scale data remain out of reach.

## Author contributions

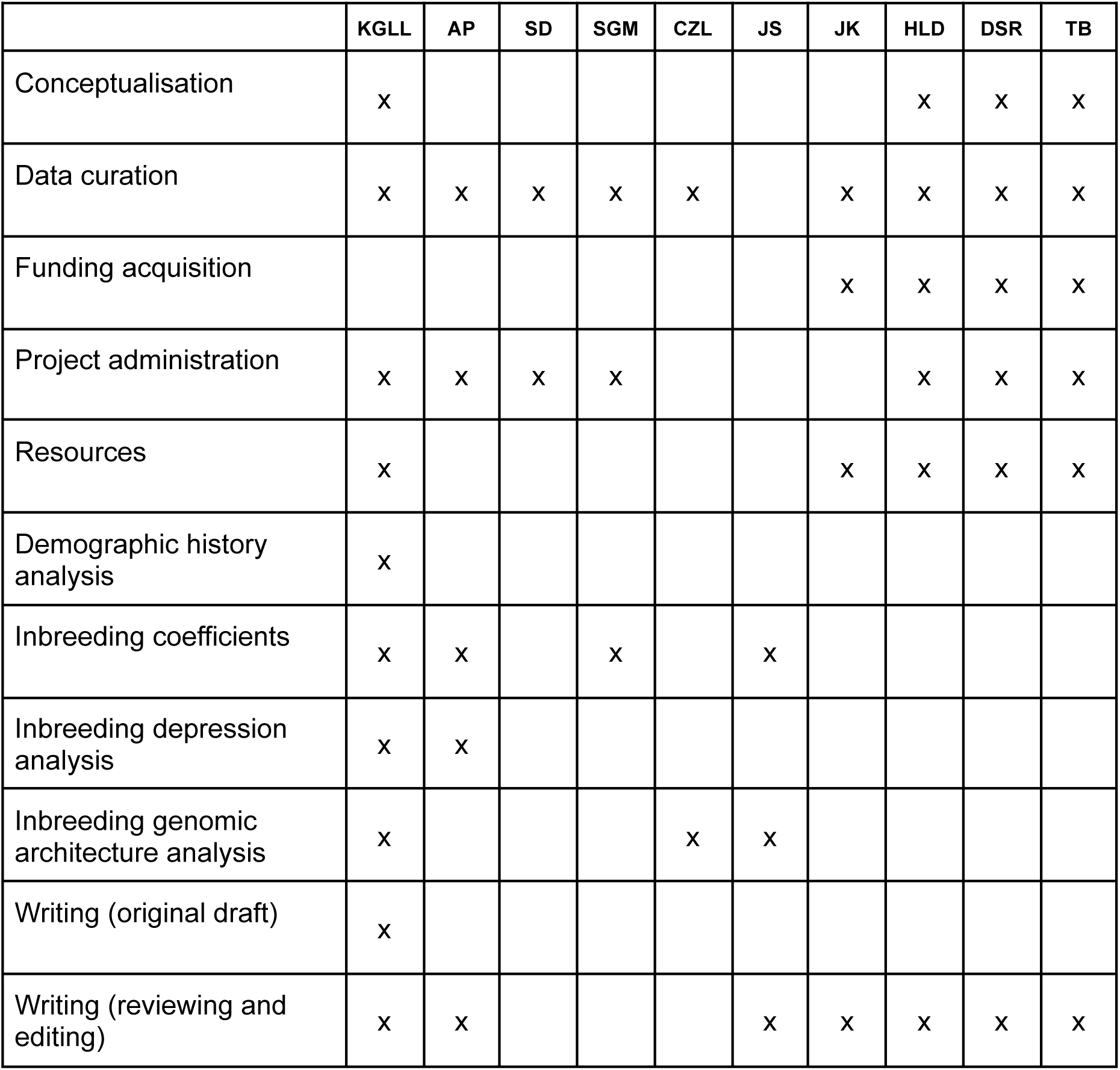

## Acknowledgements

We thank the following:

Nature Seychelles and their wardens for facilitating research on Cousin Island.

Seychelles Bureau of Standards and the Ministry of Agriculture, Climate Change and Environment for providing permission to conduct fieldwork and sample collection.

Fieldworkers, laboratory technicians, and database managers currently and previously associated with the Seychelles Warbler Project.

The IT Services at the University of Sheffield for the provision of services for High Performance Computing and troubleshooting.

The Natural Environment Research Council (NERC) grant NE/P011284/1 to H.L.D. and D.S.R for genome sequencing.

NERC grant NE/B504106/1 to T.B. and D.S.R., NWO Rubicon 825.09.013 and NERC NE/I021748/1 fellowships to H.L.D., and NWO visitors grant 040.11.232 to J.K. and H.L.D. for previous pedigree work.

NERC Environmental Omics Facility (NEOF) for sequencing and bioinformatics support. We appreciate, in particular, the technical and analytical support of Rowan Connell (University of Liverpool), Helen Hipperson, Katy Maher, Ewan Harney, Gavin Gouws, Rachel Tucker, Tom Holden and Lucy Knowles (all at the University of Sheffield).

Martin Stoffel, Tom Druet, Rebecca Shaw for their help with bioinformatics or writing scripts.

## Data availability

The genome assemblies and individual whole-genome sequence data can be found temporily at: https://zenodo.org/records/14717915 and at European Nucleotide Archive, under accession ID: PRJEB100611.

Scripts and metadata can be found at: https://github.com/kiran-lee/InbreedingDepressionSeychellesWarbler

## Supplementary material

**Figure S1.**
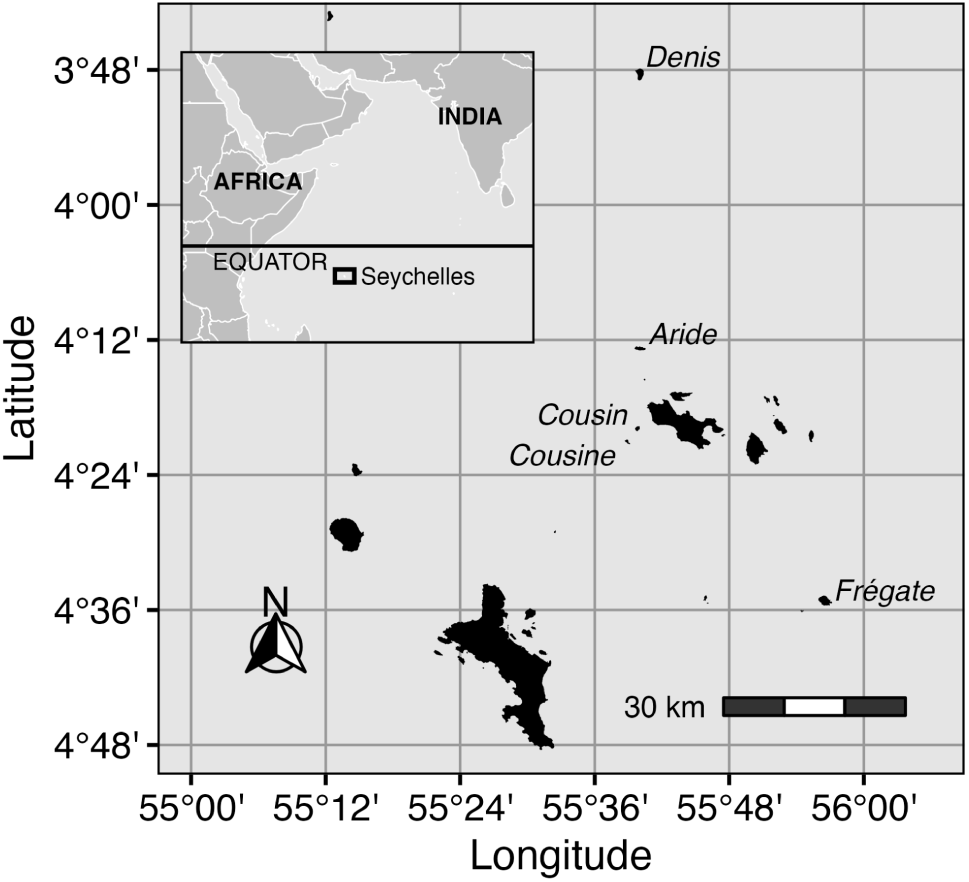
Map of contemporary Seychelles warbler populations. The global population of Seychelles warblers was bottlenecked to a remaining few on Cousin Island, which has been subject to intensive long-term monitoring. Translocations from Cousin to Aride (1988; *(Komdeur, 1994)*), Cousine (1990; *(Komdeur, 1994)*, Denis (2004; Richardson et al., 2006) and Frégate (2011; Wright et al., 2014) have established additional viable populations.

**Figure S2.**
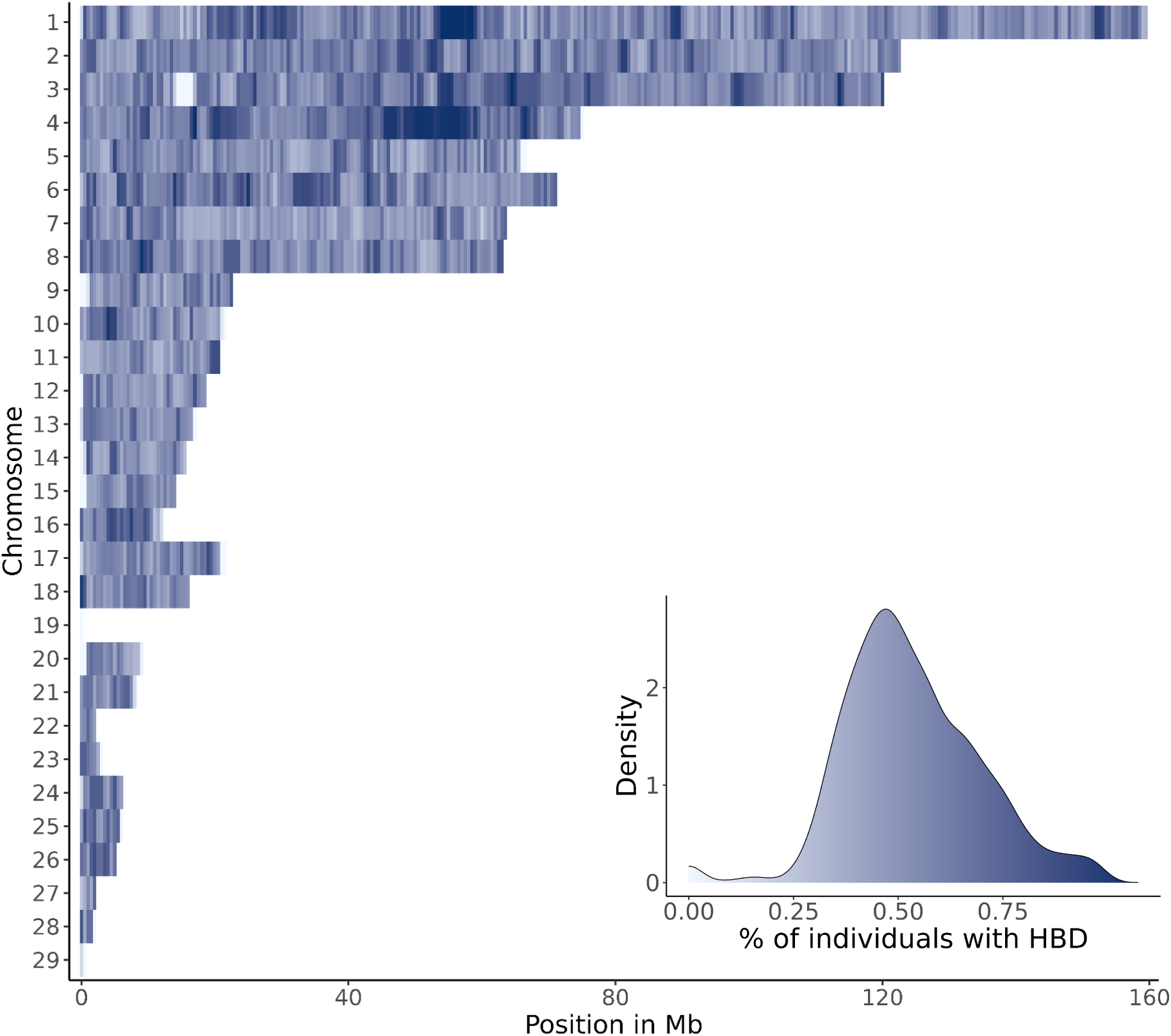
Genomic landscape of ROH in Seychelles warblers. Average ROH density per 500-kbp non-overlapping window from 1,929 individuals. Gaps in density on chromosomes 3 and 19 denote likely genome misassemblies and represent less than 1% of the total assembly.

**Figure S3.**
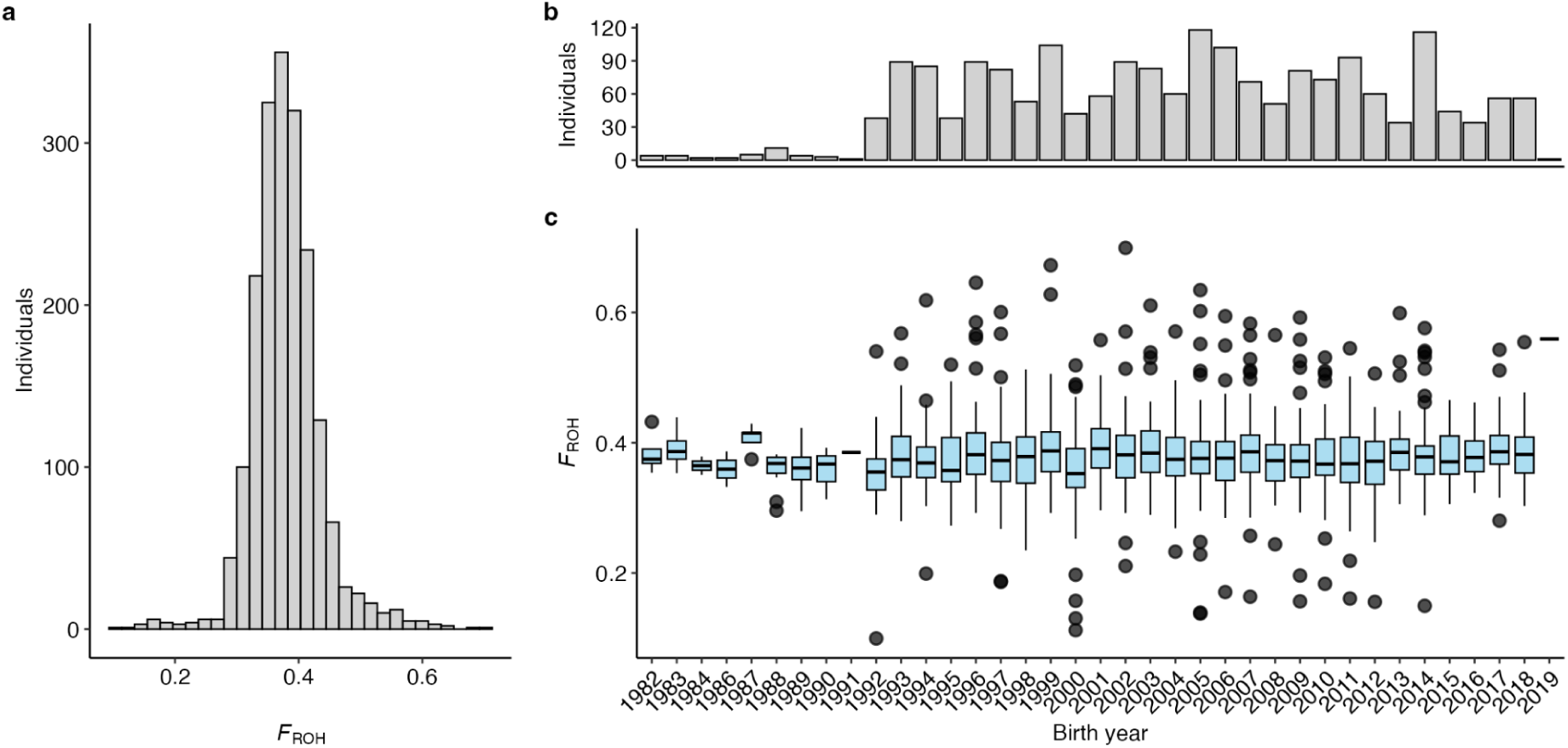
Individual inbreeding variation in Seychelles warblers. **a.** *F*_ROH_ distributions across all 1,929 individuals. **b.** Numbers of individuals sampled across annual cohorts **c.** *F*_ROH_ distributions across annual cohorts.

**Figure S4.**
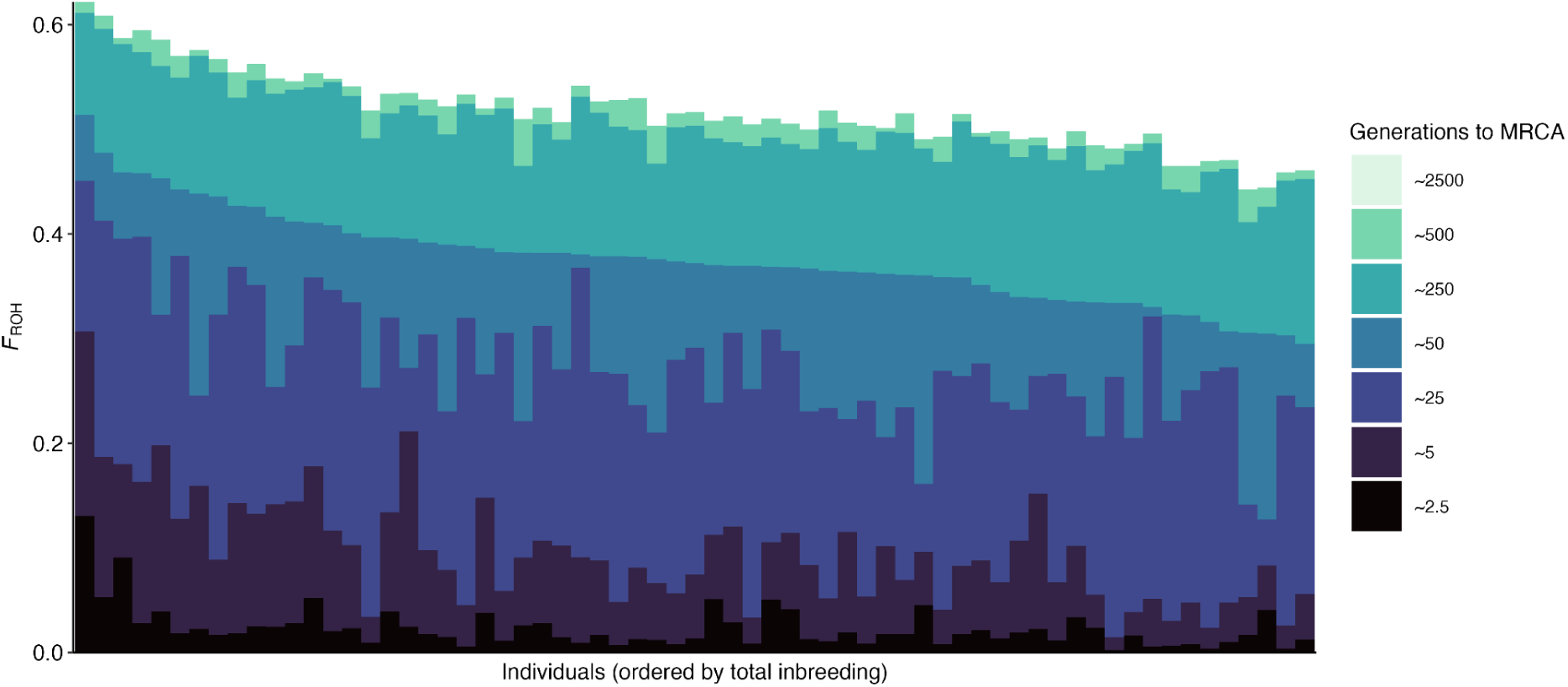
Genotyped translocation candidates. *F*_ROH_ distributions of 65 surviving genotyped individuals, ordered by recent (up to 50 generations ago) inbreeding coefficients.

**Figure S5.**
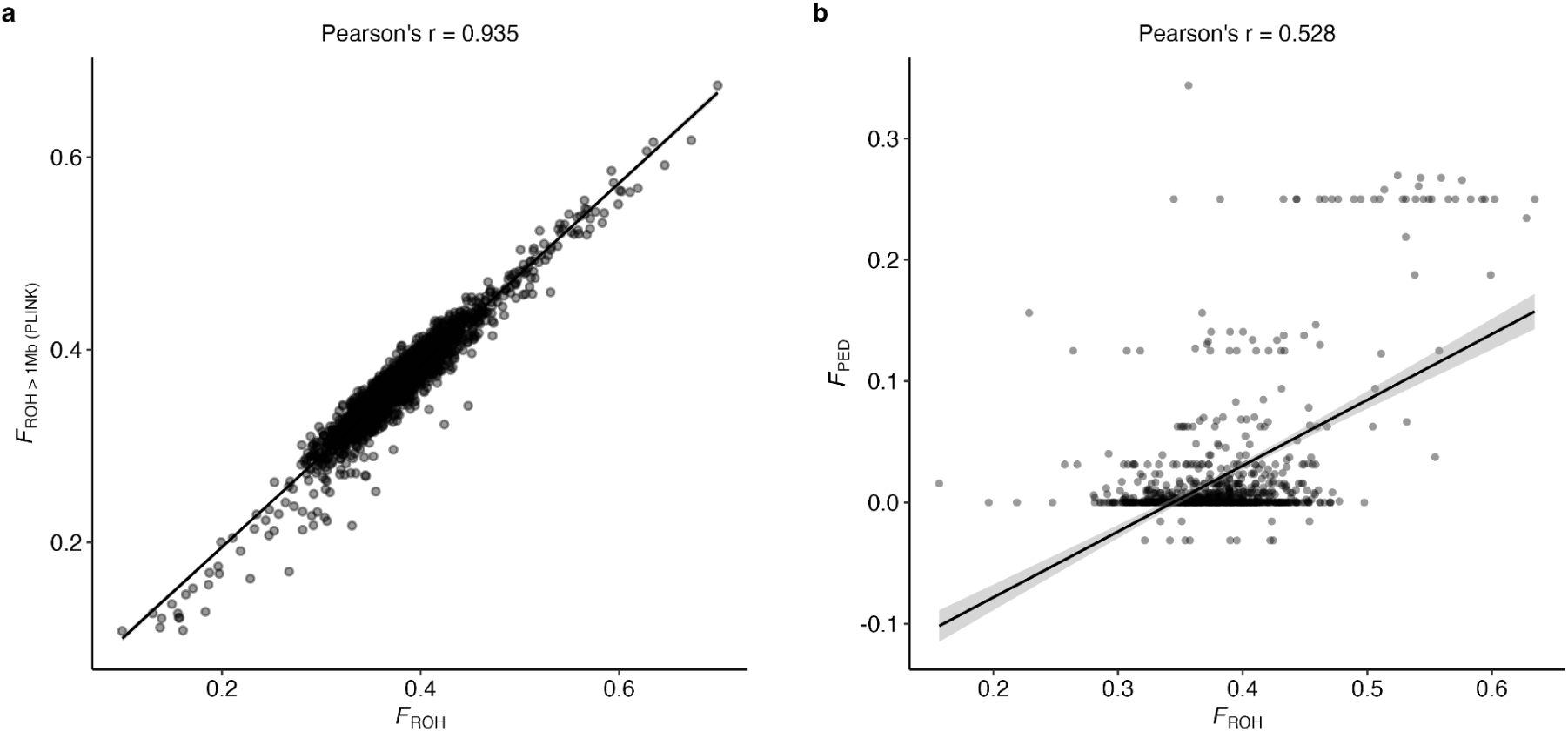
Validation of model-based inbreeding coefficient, *F*_ROH_. Correlation of HMM-derived *F*_ROH_ (RZooRoH) with **a.** rule-based approach using PLINK (n = 1,929; r = 0.93) and **b.** pedigree-based inbreeding restricted to both parents and maternal grandparents known (n = 1,104, r = 0.55).

**Figure S6.**
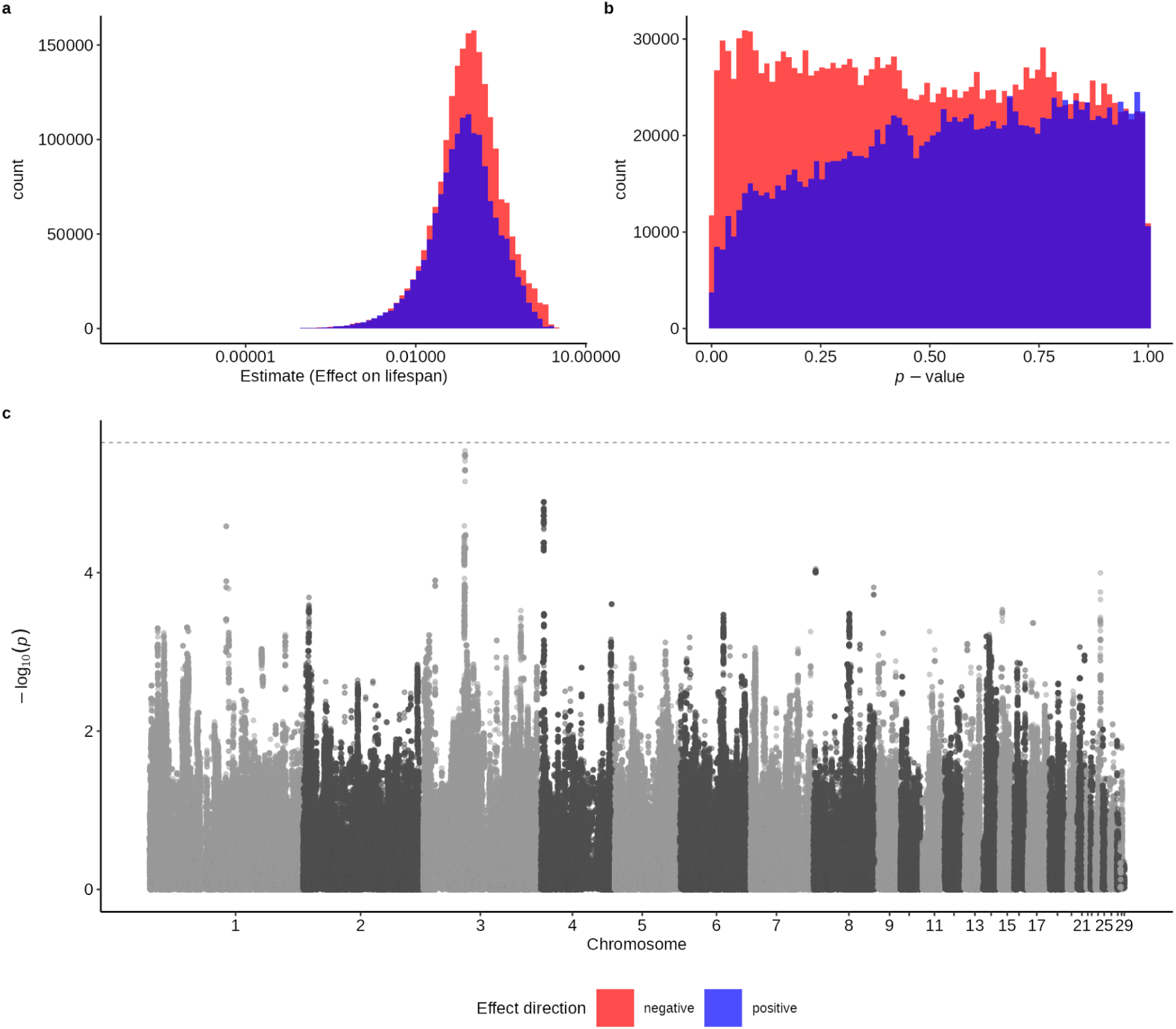
GWAS of ROH status effects on lifespan in Seychelles warblers. **a.** Distribution of effect sizes for each SNP’s allele within ROH. **b.** Distribution of *p*-values for SNPs within ROH. **c.** Manhattan plot of ROH status *p*-values across the genome. The dashed line represents the genome-wide significance threshold.

---

**Table S1.** Sex-specific summary statistics for Seychelles warbler fitness traits. Descriptive statistics for lifespan and lifetime fecundity across the whole-genome sequenced cohort (n = 1,929).

|  |  | Lifespan (years) |  |  | Lifetime fecundity (number offspring) |  |  |
| --- | --- | --- | --- | --- | --- | --- | --- |
| Sex | N | Mean | SD | Maximum | Mean | SD | Maximum |
| F | 954 | 4.69 | 4.14 | 20 | 1.74 | 2.59 | 15 |
| M | 975 | 4.44 | 3.97 | 18 | 1.75 | 3.01 | 20 |

**Table S2.** Modelling inbreeding depression in Seychelles warblers. MCMCglmm model outputs testing covariation between individual inbreeding coefficients (*F*_ROH_) and four single-generation fitness traits: **a.** annual survival, **b.** lifespan, **c.** annual fecundity and **d.** lLifetime fecundity.

| A) Annual survival | | $n = 1,735$ | | Error distribution: threshold (probit link) | |
| --- | --- | --- | --- | --- | --- |
| Term | Posterior Mean | Lower 95% CI | Upper 95% CI | Eff. Samp. | pMCMC |
| Fixed Effect |  |  |  |  |  |
| Intercept | 1.003 | 0.661 | 1.38 | 7652 | < 0.001 |
| $F_{ROH}$ (scaled) | -0.225 | -0.336 | -0.121 | 8520 | < 0.001 |
| AnnualDeathRate | -0.647 | -0.987 | -0.324 | 8236 | < 0.001 |
| Sex (Male vs Female Ref) | -0.062 | -0.17 | 0.049 | 8290 | 0.253 |
| Age_std | -0.416 | -0.655 | -0.137 | 8565 | < 0.001 |
| Age_std2 | -0.04 | -0.083 | 0.005 | 8616 | 0.076 |
| Maternal Age (std.) | -0.056 | -0.113 | -0.005 | 9000 | 0.035 |
| $F_{ROH} \times$<br>AnnualDeathRate | 0.117 | -0.45 | 0.669 | 9000 | 0.687 |
| Random effects<br>(variances) |  |  |  |  |  |
| Additive genetic | 0.024 | 0.000 | 0.073 | 9000 |  |
| Birth Year (Vby) | 0.059 | 0.011 | 0.121 | 9000 |  |
| Bird ID (Vpe) | 0.446 | 0.000 | 0.833 | 8527 |  |

| Term | Posterior<br>Mean | Lower 95%<br>CI | Upper 95%<br>CI | Eff. Samp. | pMCMC |
| --- | --- | --- | --- | --- | --- |
| Fixed Effect |  |  |  |  |  |
| Intercept | 1.102 | 1.019 | 1.176 | 7656 | < 0.001 |
| $F_{ROH}$ (scaled) | -0.197 | -0.283 | -0.108 | 8427 | < 0.001 |
| Birth Year Death Rate | -0.083 | -0.473 | 0.290 | 8702 | 0.675 |
| Sex (Male vs Female<br>Ref) | -0.023 | -0.121 | 0.077 | 8641 | 0.648 |
| Maternal Age (std.) | -0.056 | -0.106 | -0.007 | 8243 | 0.030 |
| $F_{ROH}$ (scaled) × Birth<br>Year Death Rate | -0.580 | -1.233 | 0.089 | 7705 | 0.092 |
| Random effects<br>(variances) |  |  |  |  |  |
| Additive genetic | 0.004 | 0.000 | 0.015 | 7652 |  |
| Observational-level<br>effect | 0.535 | 0.000 | 0.768 | 1458 |  |

| Term | Posterior<br>Mean | Lower 95%<br>CI | Upper 95%<br>CI | Eff. Samp. | pMCMC |
| --- | --- | --- | --- | --- | --- |
| Fixed Effect |  |  |  |  |  |
| Intercept | -1.023 | -1.270 | -0.786 | 5606 | < 0.001 |
| $F_{ROH}$ (scaled) | -0.173 | -0.259 | -0.088 | 2028 | < 0.001 |
| AnnualDeathRate | -1.620 | -3.410 | -0.118 | 11831 | 0.067 |
| Sex (Male vs Female<br>Ref) | 0.097 | -0.0103 | 0.192 | 2085 | 0.043 |
| Age_std | 1.141 | 1.069 | 1.206 | 1413 | < 0.001 |
| Age_std2 | -0.602 | -0.646 | -0.557 | 1164 | < 0.001 |
| Maternal Age (std.) | -0.006 | -0.050 | 0.041 | 2694 | 0.779 |
| $F_{ROH} \times$<br>AnnualDeathRate | 0.085 | -0.435 | 0.630 | 1957 | 0.726 |
| Variance Component<br>(G) |  |  |  |  |  |
| Additive genetic | 0.012 | 0.000 | 0.037 | 1362 |  |
| Birth Year (Vby) | 0.395 | 0.171 | 0.680 | 1197 |  |
| Bird ID (Vpe) | 0.124 | 0.077 | 0.175 | 966 |  |

| Term | Posterior<br>Mean | Lower 95%<br>CI | Upper 95%<br>CI | Eff. Samp. | pMCMC |
| --- | --- | --- | --- | --- | --- |
| Fixed Effect |  |  |  |  |  |
| Intercept | 1.091 | 1.004 | 1.180 | 9162 | < 0.001 |
| $F_{ROH}$ (scaled) | -0.163 | -0.271 | -0.056 | 9500 | 0.004 |
| Birth Year Death Rate | -0.007 | -0.439 | 0.436 | 9076 | 0.976 |
| Sex (Male vs Female<br>Ref) | 0.057 | -0.058 | 0.176 | 9500 | 0.342 |
| Maternal Age (std.) | -0.036 | -0.093 | 0.022 | 9500 | 0.223 |
| $F_{ROH}$ (scaled) × Birth<br>Year Death Rate | -0.614 | -1.356 | 0.189 | 9500 | 0.122 |
| Random effects<br>(variances) |  |  |  |  |  |
| Additive genetic | 0.003 | 0.000 | 0.013 | 9198 |  |
| Observational-level<br>effect | 0.237 | 0.000 | 0.368 | 3657 |  |

**Table S3.**
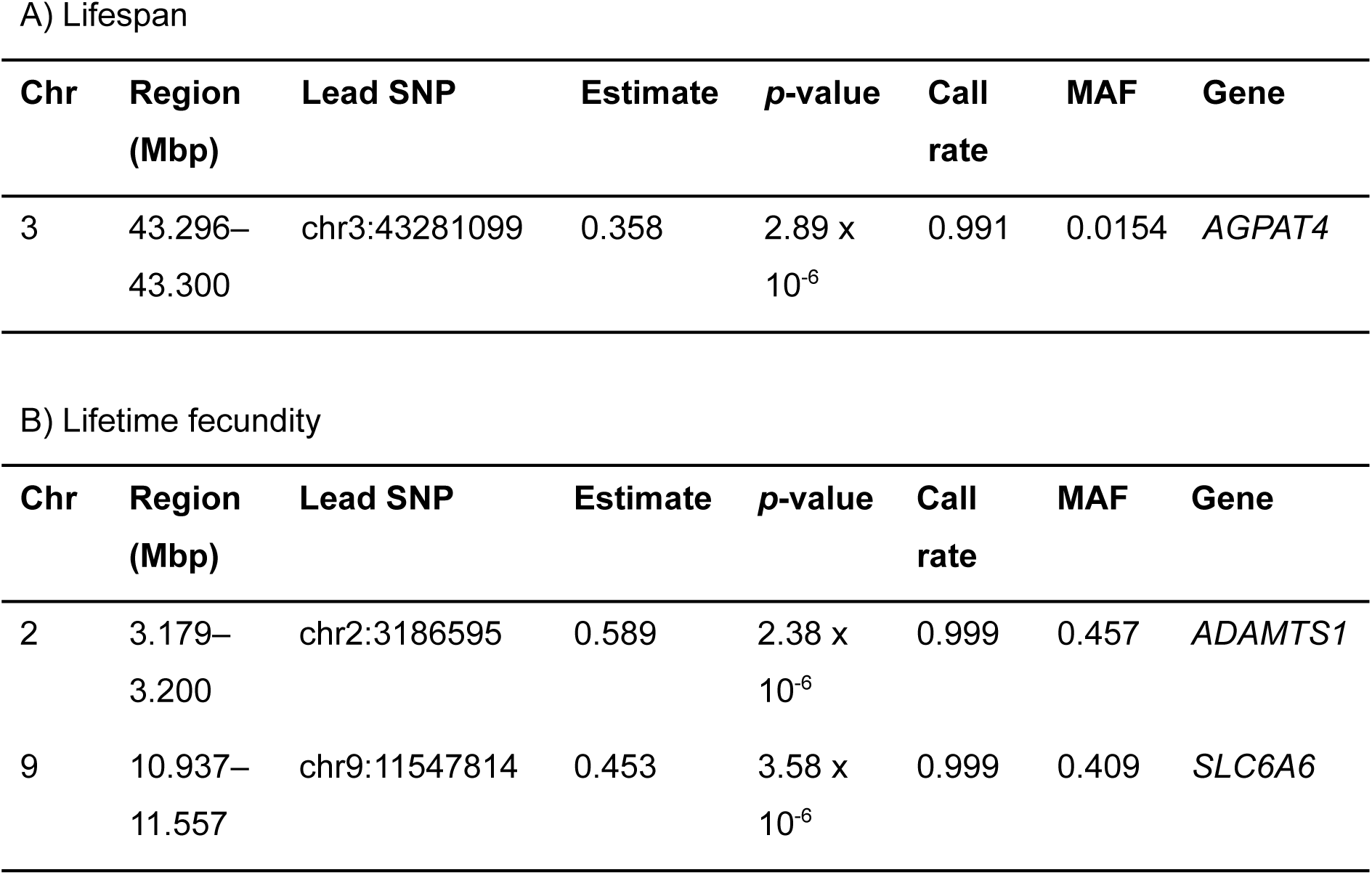
Genomic regions associated with inbreeding depression. SNP-regions of nearest significance for association between ROH status and **a.** lifespan and **b.** lifetime fecundity in the Seychelles warbler.

| Chr | Region (Mbp) | Lead SNP | Estimate | *p*-value | Call rate | MAF | Gene |
| --- | --- | --- | --- | --- | --- | --- | --- |
| 3 | 43.296–<br>43.300 | chr3:43281099 | 0.358 | 2.89 x<br>10<sup>-6</sup> | 0.991 | 0.0154 | *AGPAT4* |

B) Lifetime fecundity
| Chr | Region (Mbp) | Lead SNP | Estimate | *p*-value | Call rate | MAF | Gene |
| --- | --- | --- | --- | --- | --- | --- | --- |
| 2 | 3.179–<br>3.200 | chr2:3186595 | 0.589 | 2.38 x<br>10<sup>-6</sup> | 0.999 | 0.457 | *ADAMTS1* |
| 9 | 10.937–<br>11.557 | chr9:11547814 | 0.453 | 3.58 x<br>10<sup>-6</sup> | 0.999 | 0.409 | *SLC6A6* |

